# Mechanism of structure-specific DNA binding by the FANCM branchpoint translocase

**DOI:** 10.1101/2024.05.23.595611

**Authors:** Lara Abbouche, Vincent J Murphy, Jixuan Gao, Sylvie van Twest, Alexander P Sobinoff, Karen M Auweiler, Hilda A Pickett, Rohan Bythell-Douglas, Andrew J Deans

## Abstract

FANCM is a DNA repair protein that recognizes stalled replication forks, and recruits downstream repair factors. FANCM activity is also essential for the survival of cancer cells that utilize the Alternative Lengthening of Telomeres (ALT) mechanism. FANCM efficiently recognizes stalled replication forks in the genome or at telomeres through its strong affinity for branched DNA structures. In this study, we demonstrate that the N-terminal translocase domain drives this specific branched DNA recognition. The Hel2i subdomain within the translocase is crucial for effective substrate engagement and couples DNA binding to catalytic ATP-dependent branch migration. Removal of Hel2i or mutation of key DNA-binding residues within this domain diminished FANCM’s affinity for junction DNA and abolished branch migration activity. Importantly, these mutant FANCM variants failed to rescue the cell cycle arrest, telomere-associated replication stress, or lethality of ALT-positive cancer cells depleted of endogenous FANCM. Our results reveal the Hel2i domain is key for FANCM to properly engage DNA substrates, and therefore plays an essential role in its tumour-suppressive functions by restraining the hyperactivation of the ALT pathway.

## Introduction

Individuals carrying heterozygous or homozygous FANCM mutations are predisposed to early-onset cancer and are highly susceptible to chemotherapy-induced myelosuppression(4-7). This is because FANCM is an important mediator of DNA repair, a cellular process required to suppress cancer-causing mutations as well as respond to chemotherapy-induced DNA damage(8).

FANCM deficient cells accumulate stalled replication forks, single-strand DNA gaps, and sister chromatid exchanges, which are elevated after treatment with DNA damaging agents(9,10). Conversely, FANCM deficiency is deleterious to cancer cells which use a recombination-based mechanism of telomere maintenance (known as alternative lengthening of telomeres, or ALT). We previously showed that FANCM knockdown in ALT-positive cancer cells causes both induction of extremely high levels of replication stress and persistent recombination intermediates specifically at telomeres, resulting in cell death.(11,12) As such, FANCM is a proposed therapeutic target in ALT-positive cancers. (13,14)

FANCM performs two major functions that are critical to its role in maintenance of genome stability and ALT. First, it simultaneously binds to branched DNA molecules and translocates along the DNA(15,16). This ATP-dependent translocation activity moves the branchpoint, leading to remodeling or resolution of structures including D-loops(17), Holliday junctions(16), and R-loops(18). FANCM translocation at replication forks can also result in formation of four-way junctions, in a process known as replication fork-reversal(16). Second, FANCM acts as a molecular “anchor” for several additional DNA repair complexes on branched DNA, including the Fanconi anemia (FA) core complex and the Bloom Syndrome (BS) complex(10). Both complexes fail to localize correctly at stalled DNA replication forks in FANCM-deficient cells(10,19).

In the absence of FANCM, branched DNA structures persist and can subsequently be cleaved by structure-specific nucleases. For example, unprocessed R-loops are cleaved by XPF-ERCC1 nuclease(20), and unprocessed four-way and Holliday junctions are likely cleaved by GEN1 and the SLX4:MUS81:XPF (SMX) tri-nuclease (21). This cleavage causes the formation of double-strand breaks (9), which are the driver of increased recombination in FANCM-deficient humans or *Fancm*-knockout mice (8,22), and cell death in ALT-positive cancers after FANCM depletion (11,12). Despite a critical role for branchpoint binding in FANCM function, the mechanism by which FANCM engages branched DNA has yet to be investigated.

Other related branchpoint translocases, such as SMARCAL1, ZRANB3, and HLTF, have evolved substrate specificity domains (SSDs) to recognize branched DNA(23,24). Here, we identify and characterize a SSD in FANCM that is required to protect bound junctions from the cleavage activity of structure specific nucleases. We identify critical residues that couple structure specific DNA binding and ATPase activation, and show how FANCM binding blocks access of structure specific nucleases to junctions. Our data uncover a biochemical mechanism of FANCM binding to junction DNA and suggest dominant inhibition of FANCM ATPase activity as a therapeutic intervention in some cancers.

## Results

### The translocase domain of FANCM binds Holliday junction DNA, drives ATPase stimulation, and can perform branchpoint migration

Previous studies have shown that FANCM constructs containing the translocase domain can specifically bind branched DNA (residues 1-754) (25), resolve R-loops (1-800) (26) and have stimulated ATPase activity specifically in response to branched DNA (1-669) (27) (Figure 1A). While the translocase domain clearly enables FANCM’s branch migration activity, the specific features that drive its high affinity for branched substrates remain unknown.

**Figure 1.**
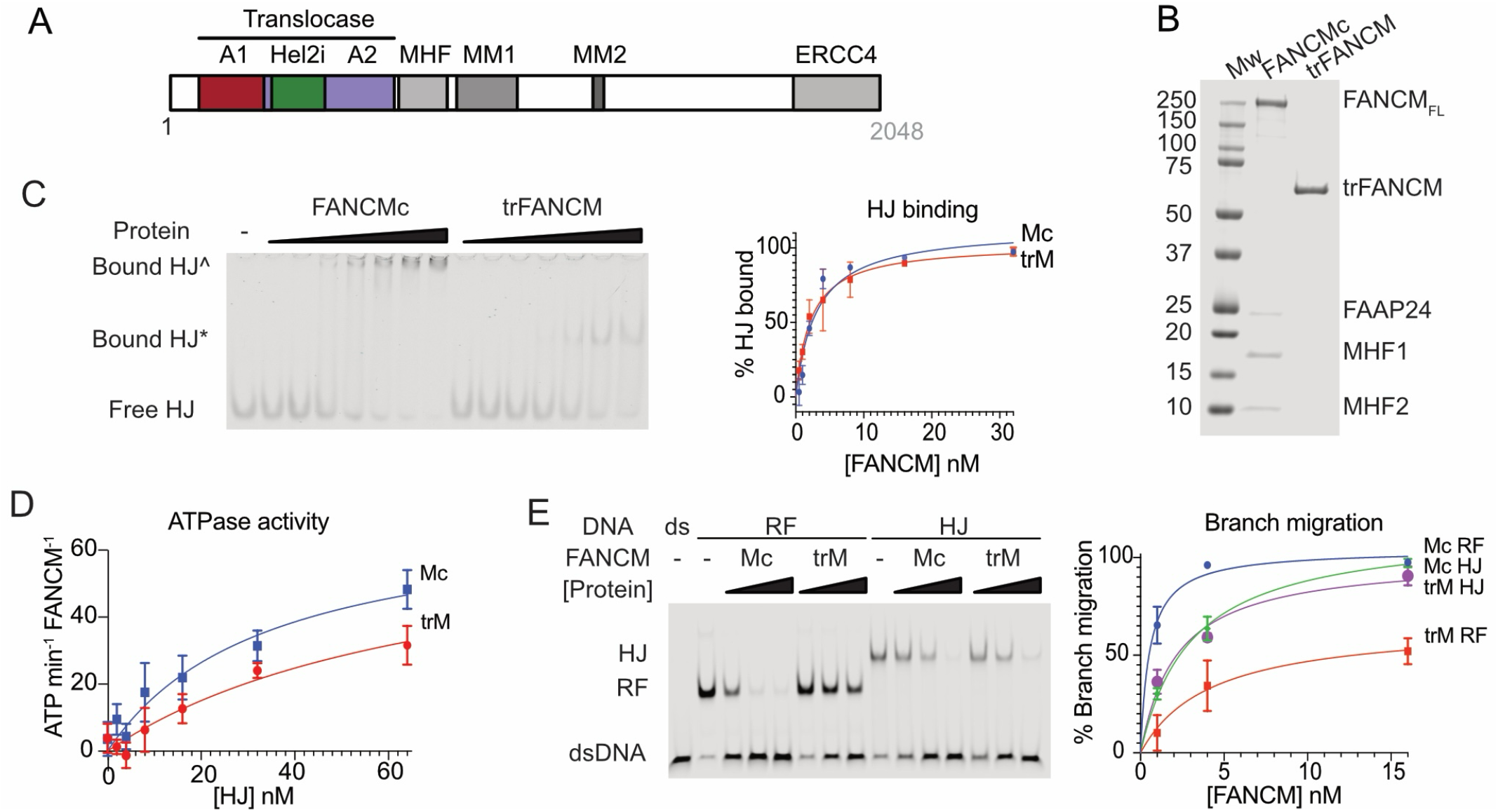
Translocase domain vs Full-length FANCM complex activity. A) Overall domain structure of FANCM showing translocase domain B) Coomassie blue stained SDS-PAGE of purified full length FANCM complex or FANCM translocase (aa82-647) C) Electromobility shift assay using oligonucleotide Holliday Junction (HJ) structures. ^=HJ bound by FANCMcFL-FANCM, *=HJ bound by FANCM_62-847_, with graph of quantification. D) Quantification of HJ-stimulated ATPase activity of FANCMc or FANCM_62-847._ E) Branch migration assays showing conversion of 3-way (RF) and 4-way (HJ) DNA structures into dsDNA, with graph of quantification.

To address this, we designed a FANCM construct (trFANCM) with just the conserved translocase domain (residues 82 to 647) with an N-terminal FLAG-tag. We expressed this protein in insect cells and purified it along with full-length FANCM complex (aka FANCMc, containing MHF1, MHF2 and FAAP24) (Figure 1B). Remarkably, the translocase domain alone displayed comparable Holliday junction (HJ) DNA binding affinity (Figure 1C) and HJ-stimulated ATPase activity to FANCMc (Figure 1D). Furthermore, it performed HJ branch migration at rates identical to FANCMc, though it was slightly less efficient at migrating replication forks (Figure 1E). Together, these data indicate that the translocase domain harbors all the necessary elements for specifically binding to and branch migrating HJ substrates.

### FANCM structural modelling predicts the Hel2i domain is involved in branch migration

To gain clues into how the translocase domain might maintain structure specificity, we generated a structural model using SWISSMODEL, vertebrate FANCM sequences and the archaeal HEF protein structure as a sequence threading template(1). The model generated by this method forms a “C” shape with two adjacent RecA-like domains (Hel1 and Hel2) separated by an insert domain (Hel2i) (Figure 2A). For confirmation, we compared it to one generated by AlphaFold2 using the orthologous method of *ab initio* modelling. The SWISSMODEL and AlphaFold models share the same overall fold (Figure 2B) and largely the same sequence register (Supplementary Figure 1A). However, the main region of divergence was in the Hel2i domain, with different placement of a helix (Homology model 430-443, AlphaFold2 model 407-426) and unstructured regions (Supp. Figure 1B). To resolve this discrepancy, we generated a third model of the Hel2i domain alone using *de novo* methods with TrRosetta. In the TrRosetta model, residues 407-445 are more consistent with the AlphaFold2 prediction but contain an additional helix (Figure 2B).

**Figure 2.**
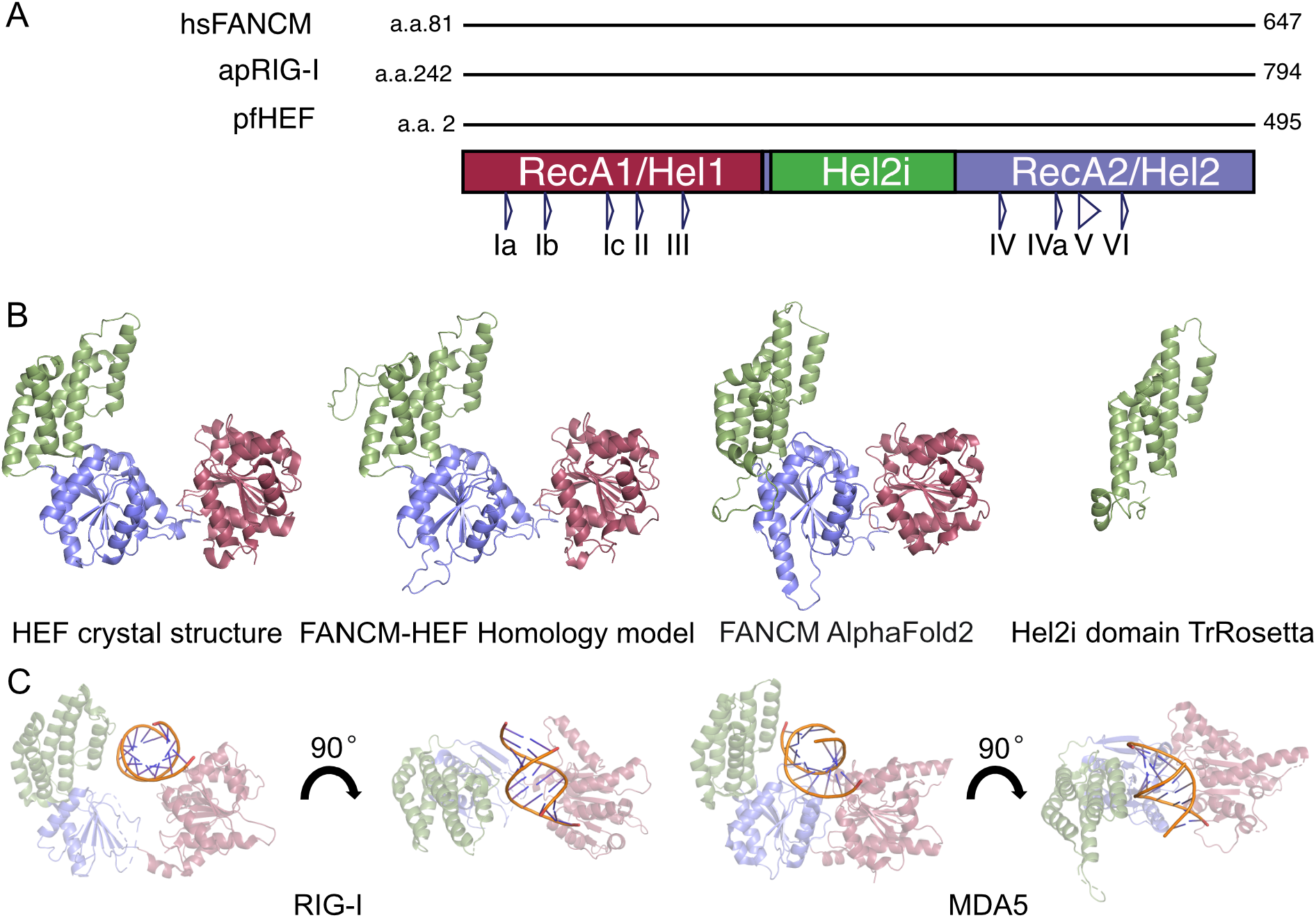
Structural models of FANCM and its comparison to evolutionarily related proteins. A. FANCM and related protein translocase domain structure, showing ATP-binding regions I-VI. B. From left to right crystal structure of HEF (1), Homology modelled FANCM, alpha fold model of FANCM, TrRosetta model of the Hel2i domain. Only residues 82-600 are shown for clarity C. Crystal structures of dsRNA bound RIG-I (PDB:2YKG ref 2) (a) and MDA5 (PDB:5JCH ref 3) showing only equivalent regions to the above FANCM models.

To predict the binding mode by which FANCM interacts with DNA, we examined structures of related proteins bound to DNA/RNA in the Protein Data Bank (PDB). Several high-resolution structures of double-stranded RNA (dsRNA) bound to RIG-I and MDA5 (see examples in (2,3,28)) are available. Although RIG-I and MDA5 are members of the FANCM protein family (29) and are involved in sensing viral dsRNA as part of the early innate immune response(2), they share structural similarities with FANCM and the HEF protein. These proteins possess the same global architecture as the FANCM models generated here, consisting of two RecA-like folds separated by an insert (Hel2i) domain.

Notably, the dsRNA is bound in a similar mode in all published structures of RIG-I and MDA5, particularly relative to the RecA1 fold, with the Hel2i domain contacting the backbone of the dsRNA and the path of the dsRNA passing over the interface between the two RecA-like folds (Figure 2C). RecA-like folds are evolutionarily ancient domains common to all SF1 and SF2 helicases that support translocation along the DNA backbone, in an ATP dependent manner. Since the major additional domain present in the translocase domain beyond the RecA-like folds is Hel2i, and this domain is in direct contact with the dsRNA in structurally related proteins, we generated the hypothesis that the insert domain in FANCM plays a key role in branch migration.

### Hel2i is the substrate selectivity domain (SSD) of the FANCM translocase domain, however other components of the FANCM complex contribute to HJ binding

To probe whether the Hel2i domain contributes to branched DNA binding, we purified FANCM complex lacking the region (residues 298-443, FANCMcΔHel2i, Figure 3A). FANCMcΔHel2i demonstrated comparable yield, stability and subunit stoichiometry as the intact complex (Figure 3B). Surprisingly, electromobility shift assays (EMSA) revealed FANCMc and FANCMcΔins to have similar HJ binding affinities (Figure 3C). Moreover, FANCMcΔHel2i retained specificity for HJs over dsDNA. By titrating in unlabelled competing dsDNA, we observed that FANCMcΔHel2i only bound branched DNA slightly less specifically than the wild-type complex (Figure 3D).

**Figure 3.**
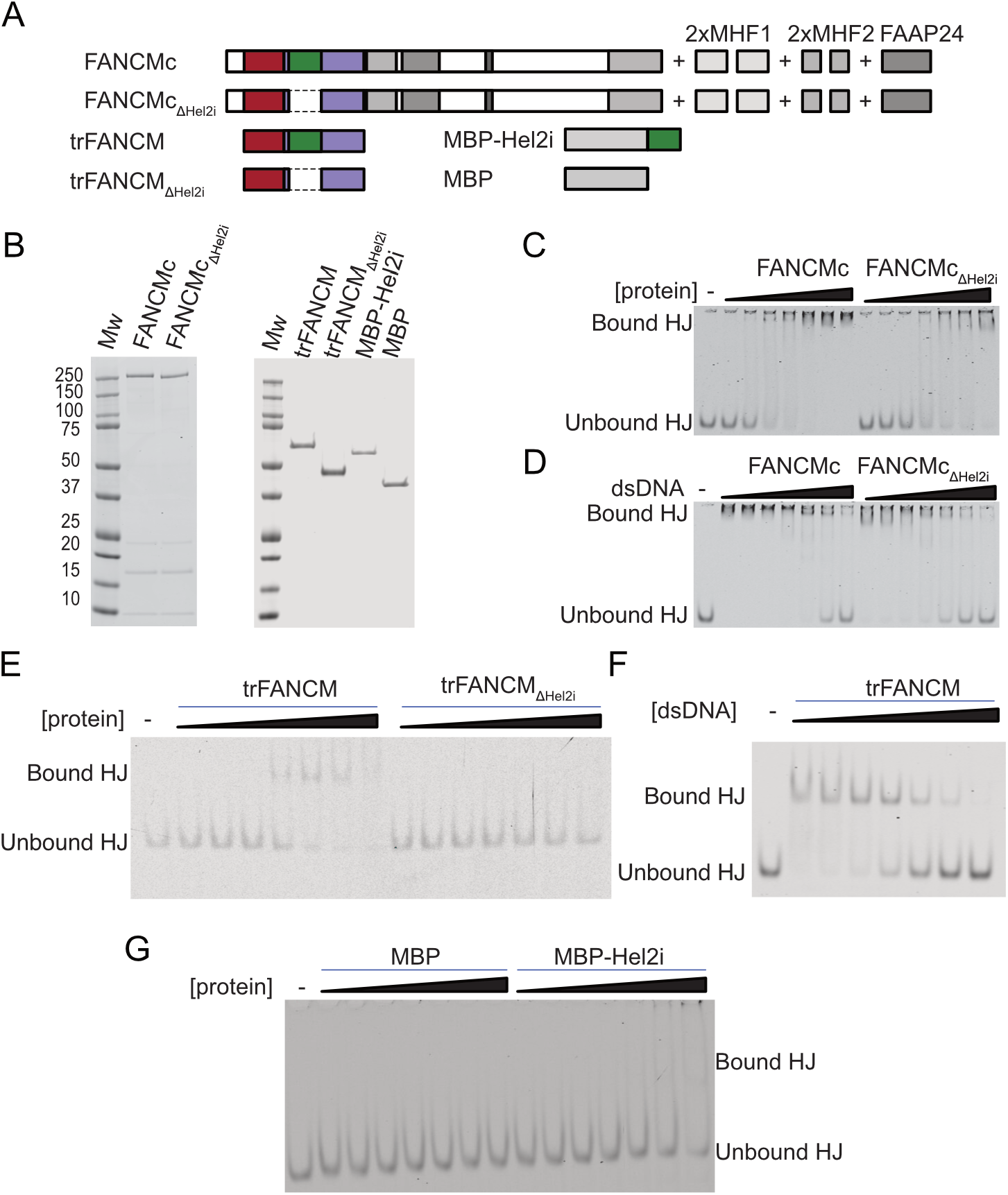
Structure specific DNA binding within the FANCM translocase domain. A) Domain structure of FANCM variants used. B) Coomassie stained WTc vs FANCMc_ΔHel2i_, trFANCM and trFANCM_ΔHel2i_, MBP-Hel2i and MBP C) Electromobility shift assays (EMSA) using recombinant FANCMc EMSA vs FANCMc_Δins_ and 4-way HJ DNA. D) HJ EMSA with competition from increasing concentrations of unlabelled dsDNA. E and F) As in C and D, but using translocase only FANCM proteins. G) EMSA of HJ using MBP-Hel2i domain or control MBP-only protein.

Since FANCM contains other structure-specific binding domains outside of the translocase domain (e.g., MHF1, MHF2, and FAAP24), we examined HJ binding by isolated constructs containing only the translocase domain (trFANCM, Figure 3A). This region showed a similar specificity for branched DNA to the full FANCM complex when measured by dsDNA competition assays. But trFANCMΔHel2i exhibited significantly lower affinity for HJ binding, with at least a 60-fold reduction in binding affinity compared to trFANCM (Figure 3F).

To determine if Hel2i only could bind DNA, we purified it in isolation, fused to an MBP solubility tag (Figure 3A-B). We found that the Hel2i domain exhibited only weak HJ binding (Figure 3F). Together with the results above, this indicates that the Hel2i domain confers HJ specificity to the N-terminal translocase domain dependent upon its positioned integration within the RecA2 domain architecture (29).

### Hel2i enables activation of FANCM ATPase activity necessary for DNA translocation

We tested FANCMcΔHel2i and trFANCMΔHel2i mutants in assays that measure FANCM branch migration activity. Compared to wild-type, these mutants exhibited a complete loss of HJ branch migration (Figures 4A-B). Furthermore, they were unable to resolve R-loops (Figure 4C), another branched DNA substrate acted upon by FANCM. To confirm that the loss of activity in trFANCM-ΔHel2i was not due to the absence of an intervening sequence, we also tested trFANCM-RIGI, a chimeric protein in which the FANCM-Hel2i domain was replaced with the RIG-I-Hel2i domain. This chimeric enzyme was also completely inactive in HJ branch migration assays (Figure 4B), despite presence of an intervening sequence.

**Figure 4.**
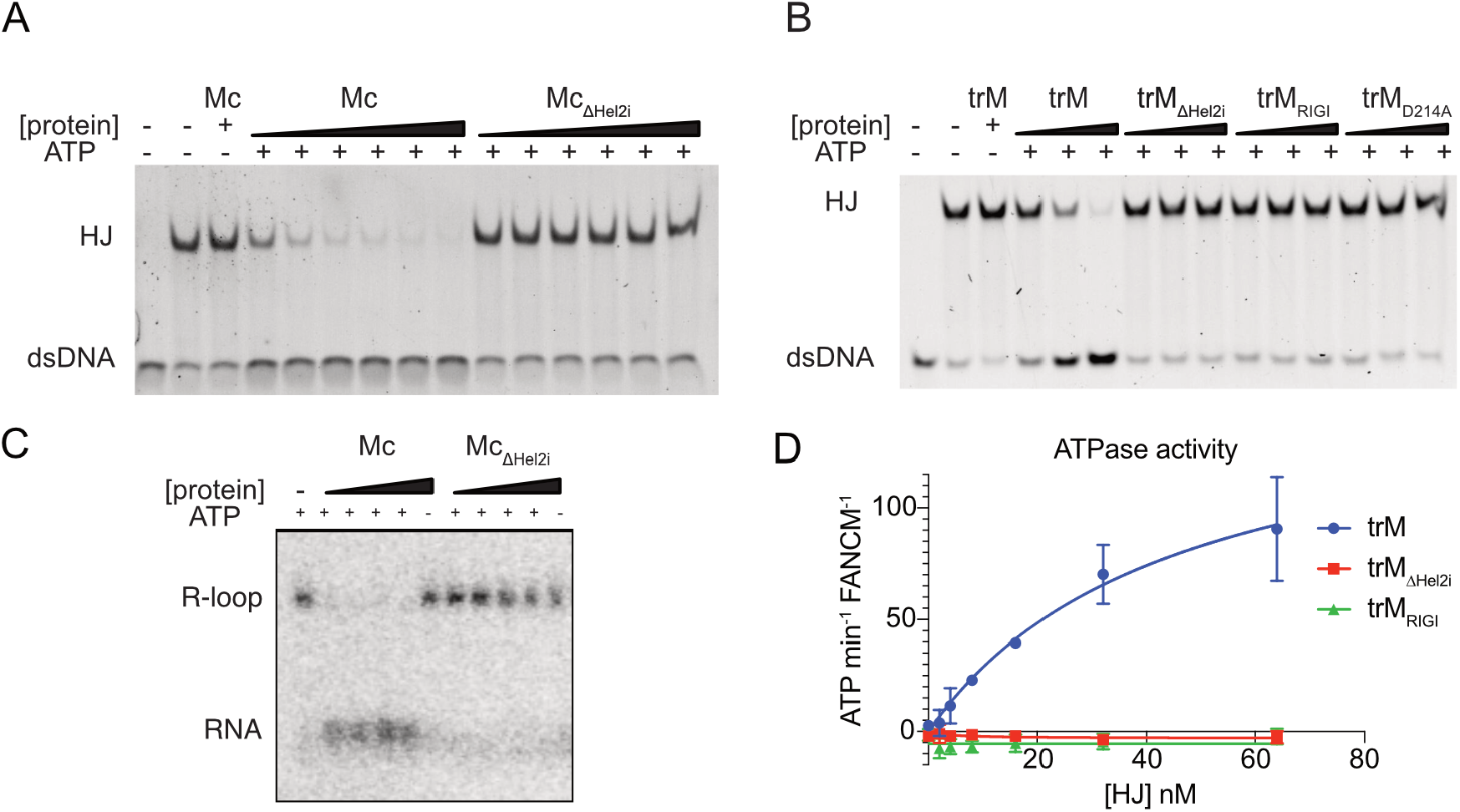
The Hel2i domain is essential for branch migration of Holliday junctions (HJ) and R-loops, dependent upon its ATPase domain. A) Branch migration activity of full length FANCMc and FANCMc_ΔHel2i_ on HJ substrate B) Branch migration activity of trFANCMc, trFANCM_RIGI_ and trFANCMc_Δins_ translocase only proteins C) R-loop resolution activity of FANCMc and FANCMc_ΔHel2i_ on 140bp R-loop structure within 2.1kb plasmid D) HJ-dependent ATPase stimulation of trFANCMc, trFANCM_RIGI_ and trFANCMc_ΔHel2i_

While the MHF1/2-containing FANCMcΔHel2i complex retains HJ binding capability (Figure 3), these results demonstrate that the Hel2i domain is indispensable for effective engagement and branch migration of branched DNA substrates like HJ and R-loops.

To investigate the DNA-dependent stimulation of FANCM’s ATPase activity, we tested multiple DNA structures for their capacity to activate the enzyme. Our data reveal that linear duplex DNA molecules exhibit a low capacity to activate the ATPase activity, while splayed arm structures are more efficient activators. Notably, increasing the length of double-stranded DNA adjacent to the branchpoint further stimulates the enzyme’s activity (Supplementary Figure 2).

Using the 15bp Holliday junction, we investigated DNA-stimulated ATPase activity in the trFANCMΔHel2i and trFANCM-RIGI mutants. Remarkably, despite these mutants having intact RecA1 and RecA2 folds, which are requisite for ATPase motor activity, they exhibited a complete absence of both baseline and DNA-stimulated ATPase activity (Figure 4D). Circular dichroism experiments confirmed that these proteins are properly folded (Supplementary Figure 3), ruling out potential misfolding as a cause for the observed lack of activity. These results underscore the critical importance of the Hel2i domain for FANCM’s ATPase activity, even in the presence of branched DNA substrates. While the RecA1 and RecA2 folds are necessary for the ATPase motor function, the Hel2i domain appears to be indispensable for coupling DNA binding to ATP hydrolysis and enabling the enzyme’s robust activation upon encountering branched DNA structures.

### Key residues in the Hel2i domain of FANCM are essential for both HJ binding and ATPase activity

To identify the key residues within FANCM-Hel2i that play the specific branched DNA activation role, we generated a model of FANCM bound to a splayed DNA splay using Alphafold3. The model places the RecA folds in contact with the dsDNA and fittingly, the insert is in direct contact with the branchpoint of the splay, parting it in the middle. The position of the branchpoint is such that if two more strands were added to the splay to make a HJ – the Hel2i domain would act as a pin, inserted into the centre of the branchpoint. This inserted pin, coupled with the pushing activity of the RecA folds, parsimoniously explains the branch migration mechanism of FANCM, the specificity of FANCM for binding branched structures, the importance of the insert domain for binding branched structures and its absolute necessity for branch migration (Figure 5A). A series of targeted point mutations were selected to introduce major chemistry changes to residues predicted to mediate DNA binding along with residues predicted to not be involved in DNA binding (Figure 5B).

**Figure 5.**
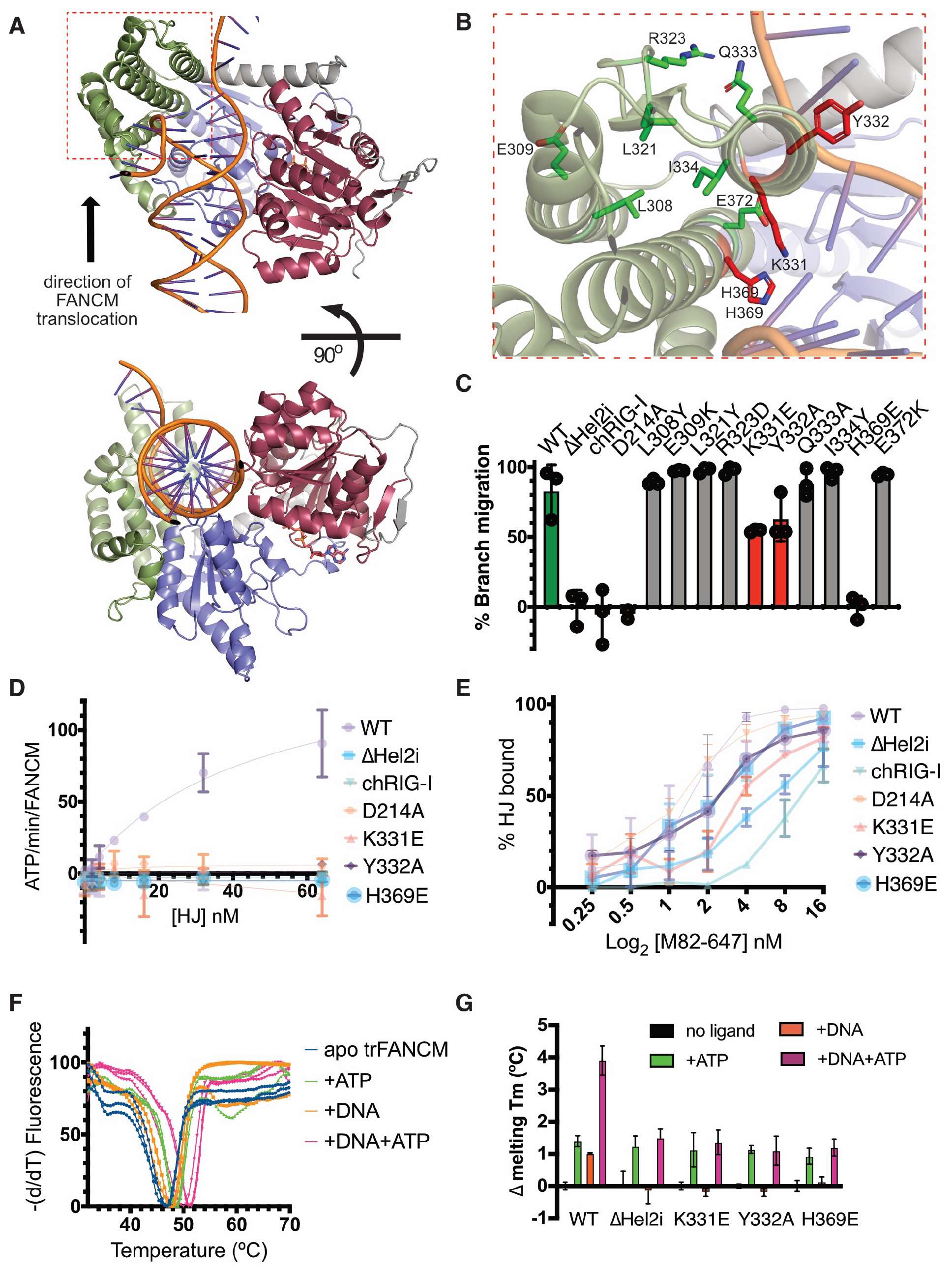
Key residues in the FANCM-Hel2i domain govern engagement with, and activity on, branched DNA,. A) AlphaFold 3 model of FANCM translocase domain engaged with a splayed DNA molecule, colourised as per Figure 2, with residues C-terminal of the RecA2 fold shown in grey. and B) Zoom in of (A) showing key Hel2i residues predicted to bind at or near junction and mutated for testing in branch migration assays. C) Effects of indicated single amino acid changes in Hel2i on HJ branch migration activity, (endpoint unwound product) D) DNA stimulated ATPase activity and E) HJ binding by EMSA for indicated mutants. F) Example DSF assays showing three replicate denaturation experiments of trFANCM in the absence or presence of ATP, DNA or DNA+ATP. G) Change in overall stability of trFANCM or indicated mutants as measured by DSF. Only WT trFANCM shows increased thermal stability upon addition of DNA, while all variants show an ATP-dependent shift.

To test our predictions, we purified the designed panel of FANCM mutants and examined their activities on synthetic HJ substrates (Figure 5C). Several mutants, including K331E, Y332A and H369E, exhibited substantially reduced HJ branch migration compared to wild-type FANCM, consistent with disrupting critical DNA contacts at the branch point. For H369E no detectable migration was observed, corresponding to that seen for trFANCMΔHel2i and trFANCM-RIGI lacking the entire Hel2i domain, or the D214A ATP-hydrolysis deficient mutant.

We next examined the impact of the K331E, Y332A, and H369E mutations on FANCM’s ATPase activity and its binding affinity to HJ (Figures 5D-E). All three inactive mutants exhibited significantly reduced HJ binding compared to the wild-type trFANCM. Consequently, they lacked detectable DNA-stimulated ATPase activity, suggesting that the mutations disrupted critical DNA contacts at the branchpoint required for coupling DNA binding to ATP hydrolysis. Collectively, these mutational analyses provide strong experimental support for our structural model, highlighting key residues in FANCM’s Hel2i domain, such as K331, Y332, and H369, that are crucial for recognizing and promoting branch migration of HJ structures.

One mechanism by which the Hel2i domain could activate FANCM’s ATPase activity upon DNA binding is through inducing a structural stabilisation of the enzyme when contacting branched DNA but not dsDNA. To test the conformational stability of trFANCM, we employed differential scanning fluorimetry (DSF), which measures the temperature-induced loss of protein stability by monitoring the binding of a fluorescent dye to exposed protein surfaces(30). ATP or branched DNA induced a reproducible stabilisation of trFANCM of 1.5ºC and 1ºC respectively, while addition of both ligands together increased the overall stability by 4ºC (Figure F). As the fluorescent signal in DSF comes purely from protein, this result confirms that the FANCM translocase domain adopts a conformational stabilisation when bound to DNA, that is further stabilised when it is in an ATP-bound form.

The melting temperature of trFANCM-ΔHel2i differed slightly to trFANCM, likely due to the size and shape-dependent effects of DSF (30), but still had a 1.5 ºC ATP-induced shift in melting temperature (Figure 5F). However, the other Hel2i mutant versions, being of identical size, displayed similar melting temperatures in the apo and ATP bound forms (Figure 4G). In contrast, the ΔHel2i and Hel2i point mutants showed no further stabilisation upon addition of DNA. These findings confirm that while the Hel2i mutant proteins retain the capacity to bind ATP, they fail to bind DNA. This observation aligns with previously described substrate- and ATP-dependent conformational changes occurring in RIG-I and MDA-5 translocases (28,31), suggesting that the Hel2i domain is crucial for coupling DNA binding to the ATP-dependent conformational changes in FANCM.

### FANCM binds an open HJ conformation and protects from cleavage by structure specific nucleases

Due to the flexible nature of DNA and the crossover point, HJs can adopt multiple conformations: an open form with a four-way symmetric X-structure, and two stacked forms where pairs of arms are aligned in a parallel or anti-parallel manner(32) (Figure 6A). While the anti-parallel form is observed in X-ray crystal structures of naked HJs, its existence *in vivo* is likely prevented by physical constraints imposed by the flanking arms(2). To determine the conformation of the HJ bound by FANCM, synthetic BXRH Holliday junctions were used. These junctions have four arms (B, X, R, and H) with varying lengths, affecting the overall DNA shape: in the absence of cations like Mg^2+^, the negative charge of the DNA backbone favours an open conformation while in the presence of cations they adopt a parallel stacked conformation (33). The addition of Mg^2+^ altered the mobility of the BXRH junctions from predominantly open to predominantly closed (stacked) conformation, which is measured by their altered mobility in polyacrylamide gels. Importantly, this stacked conformation prevented the binding of FANCM, suggesting that FANCM favours binding to only the open form of the HJ. Under these *in vitro* conditions, the translocase domain alone cannot convert the stacked conformation into the open form in the presence of physiological levels of Mg^2+^.

**Figure 6.**
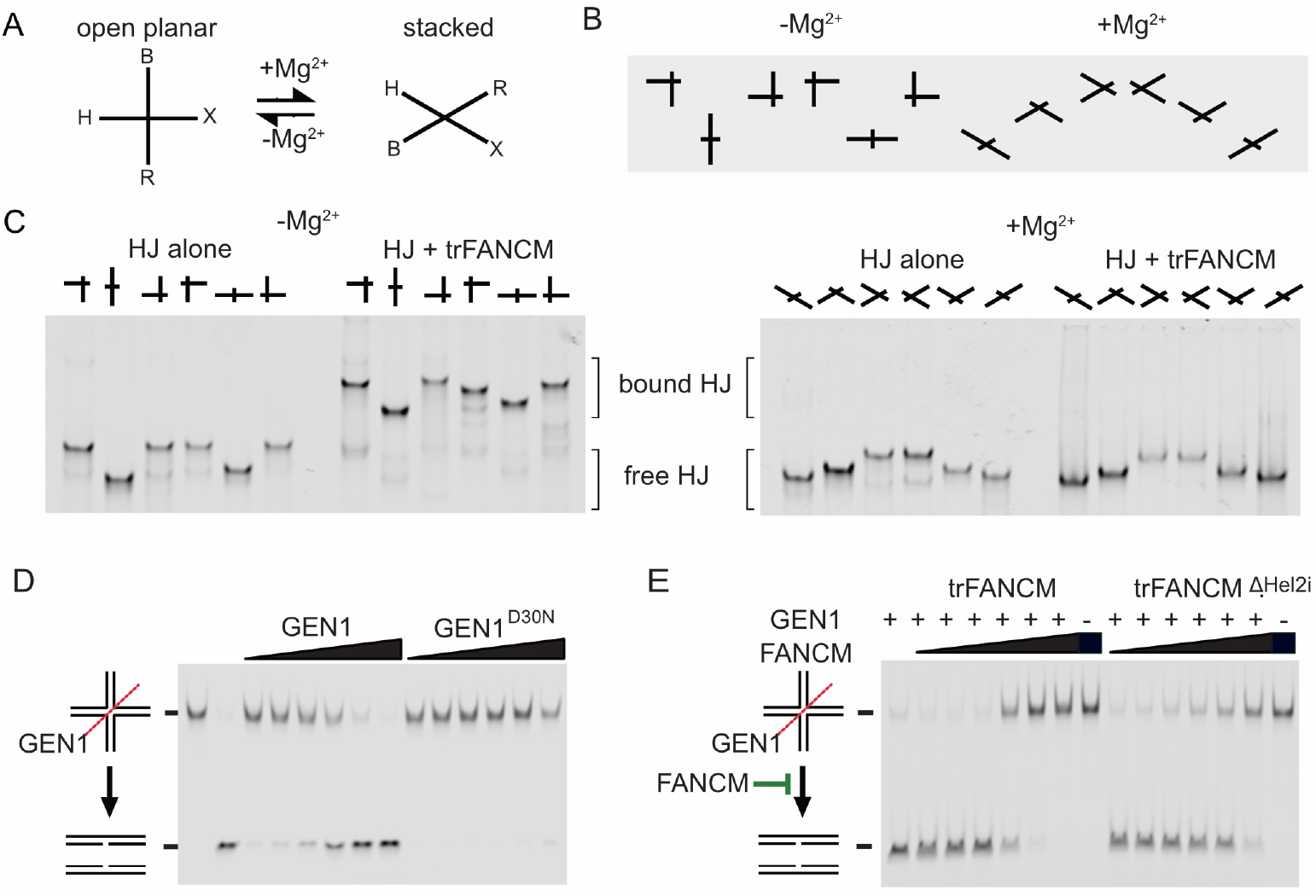
FANCM binds the open form of HJ. Schematics of A) open planar and stacked forms of HJ and B) the set of 6 BXRH HJ of different arm lengths, showing their relative mobility in polyacrylamide gels. C) Electromobility shift assay (EMSA) of BXRH HJ substrates with and without trFANCM in the absence of presence of 1mM MgCl_2_ in the sample and running buffer. D) Cleavage of 10nM static HJ by 1, 3, 10, 30, 90 or 270nM GEN1 or nuclease dead GEN1-^D30N^. E) Cleavage of HJ by GEN1 (4nM) in presence of 12.5, 25, 50, 100, 200, or 400nM trFANCM or trFANCM^ΔHel2i^

In addition to branch migration, HJ can also be cut by HJ resolution enzymes called resolvases, which can cleave HJ to generate crossover and non-crossover products during homologous recombination (34). In bacterial recombination, RuvC cleavage activity has been shown to be stimulated by RuvAB translocase binding, which, like FANCM, stabilizes the open HJ structure (35). This led us to hypothesise that FANCM may play a similar stimulatory role on HJ cleavage. To test this, we performed GEN1-mediated HJ cleavage assays with and without FANCM.

As previously published, cleavage of labelled HJ by GEN1 converts it into two linear, faster migrating DNA products (Figure 6D). Strikingly, the presence of FANCM at concentrations equimolar with GEN1 caused partial inhibition of HJ cleavage, and at 10x enzyme ratios caused complete inhibition (Figure 6D). Increasing concentrations of trFANCMΔHel2i only partially blocked GEN1 activity at 50x excess concentrations (Figure 6E). These findings led us to conclude that trFANCM binds at the junction point and blocks GEN1 action, even though the junction is stabilised in the preferred conformation for cleavage. As GEN1 is a nuclease that could potentially cleave the junction any time it becomes unbound by FANCM, the high potency of FANCM inhibition suggests a stable interaction with a relatively long half-life. Unlike in bacterial systems, translocase binding prevents access by other HJ processing enzymes, leading to protection of the HJ structure.

### Integrity of the FANCM Hel2i is required for viability of cells utilising the ALT pathway of telomere maintenance

Our previous studies demonstrated that FANCM knockdown hyperactivates hallmarks of the Alternative Lengthening of Telomeres (ALT) mechanism, including increased telomeric single-stranded DNA (ssDNA) production and reduced break-induced telomere synthesis(11-13). This activity required FANCM’s interaction with the Bloom syndrome complex but not the Fanconi anaemia core complex, indicating that FANCM curtails excessive ALT activity by alleviating recombination-mediated replication stress at telomeres. Consequently, several groups have reported that FANCM deletion or inhibition can selectively kill ALT-positive cells(11,12,36). To determine whether hyperactivation of ALT phenotypes occurs in the absence of structure-specific DNA binding by FANCM, we expressed wildtype or Hel2i-mutant FANCM in the ALT-positive U2OS cell line, along with simultaneous knockdown of endogenous FANCM (Figure 7A-B). Knockdown of FANCM in U2OS cells resulted in a robust accumulation of telomeric single-stranded DNA (ssDNA) marked by phosphorylated RPA, indicative of telomere-specific replication stress, and an accumulation of cells in the G2/M phase of the cell cycle (Figure 7C-D). Complementation with wildtype FANCM largely restored low ssDNA levels and normal cell cycle progression. However, the DNA binding-deficient FANCM mutants, FANCM-ΔHel2i or FANCM-H369E, failed to rescue these phenotypes (Figure 7C-D), similar to the ATPase-defective mutant FANCM-K117R. Double mutants harbouring both DNA binding and ATPase deficiencies exhibited similar effects as the single mutants.

**Figure 7:**
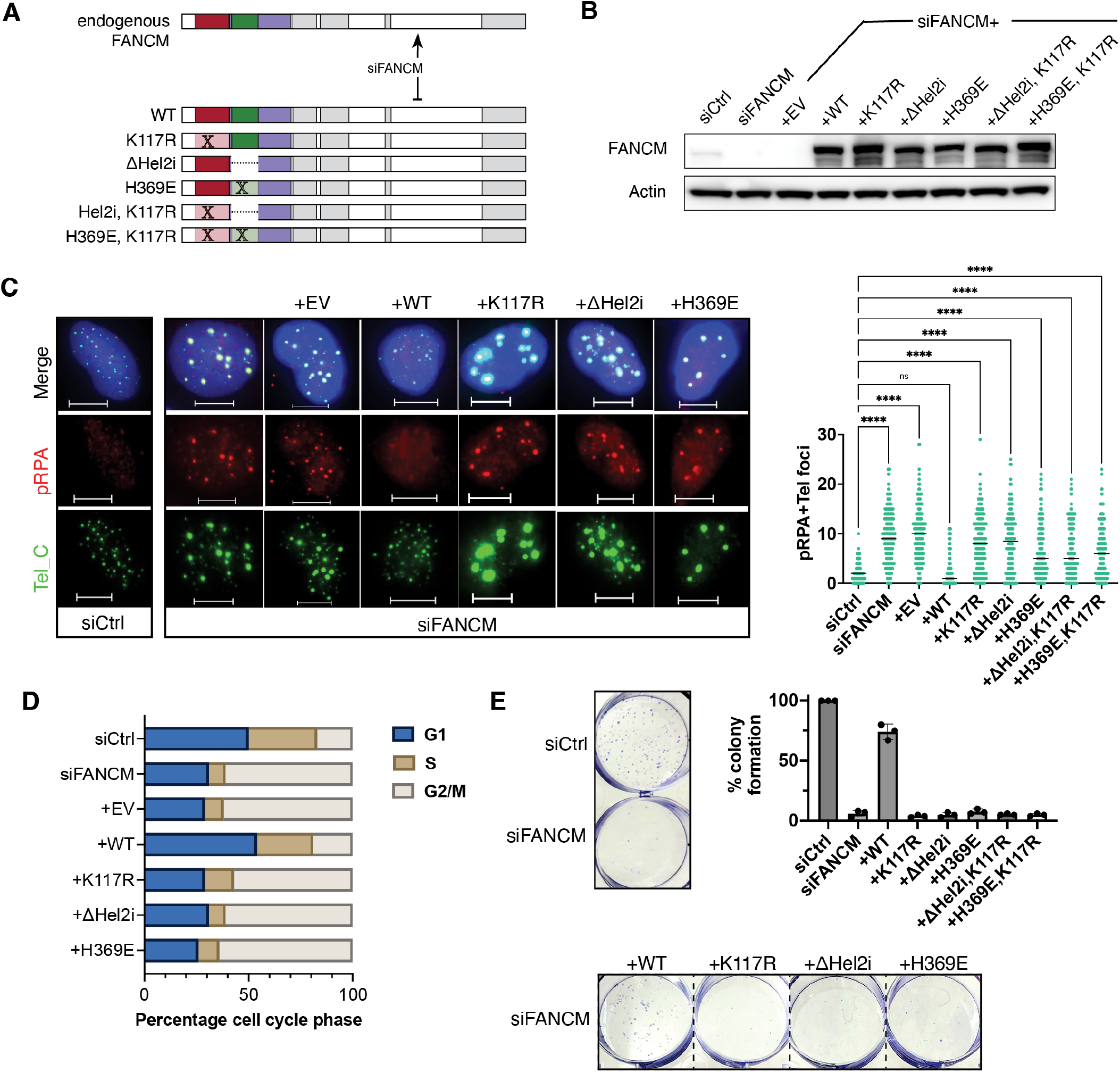
FANCM Hel2i domain integrity is essential for the survival of ALT+ U2OS cells. A) siFANCM targets endogenous but not exogenous FANCM mRNA for degradation in U2OS cells B) Western blot showing knockdown and re-expression of FANCM wildtype or mutants. C) RPA/telomere FISH co-staining in example cells (scale bar = 10μm), along with quantification of at least 100 nuclei in each line. Statistical analysis vs siCtrl by One Way ANOVA (****p<0.0001, ns=not significant) D) Quantification of cell cycle profiles as measured by propidium iodide staining and flow cytometry, E) Example clonogenic survival assay and quantification from n=3 experiments performed in triplicate.

We next measured the effects of the different FANCM variants on the viability of U2OS cells depleted of endogenous FANCM. While re-expression of siRNA-resistant wildtype FANCM significantly rescued the viability defect caused by endogenous FANCM depletion, all other mutants failed to do so, exhibiting at least 12-fold reduced viability comparable to vector-transduced controls (Figure 7E). These results demonstrate that disrupting the Hel2i-mediated branched DNA recognition impairs FANCM’s capacity to restrain hyperactivity of the Alternative Lengthening of Telomeres, making it an essential suppressor of this tumour-specific telomere maintenance mechanism.

## Discussion

FANCM is a key genome stability and tumour suppressor protein that is also essential for the viability of cancer cells that utilise Alternative Lengthening of Telomeres (ALT) for survival. Its central role involves the ability to simultaneously bind branched DNA structures and translocate along DNA, resulting in branchpoint migration (37). This allows FANCM to remodel or resolve structures such as D-loops (17), R-loops(18), Holliday junctions(16) and replication forks(17). Upon binding branchpoints, FANCM also acts as a molecular anchor for additional DNA repair complexes(10). Despite the importance of structure-specific DNA binding to its function, the mechanism by which FANCM engages and facilitates branch migration of branched DNA structures was unknown.

In the present work, we have shown that the N-terminal translocase domain of FANCM provides a minimally competent region for binding HJ and other branched DNA structures, followed by their branch migration. Within the translocase is an evolutionarily conserved region, Hel2i, inserted within the canonical RecA2/Hel2 fold. The presence of this domain is intriguing, as it is present in only a few other human translocases, all of which are involved in RNA metabolism. These include the pattern recognition receptors RIG-I, MDA5 and LGP2 involved in viral RNA recognition, and the nuclease DICER involved in generation of miRNAs and siRNAs. For RIG-I and MDA-5, where structures have been solved, the Hel2i domain directly contacts the bound substrate (2,3,28,31). Together with the RecA folds, the RNA substrates are almost entirely encircled by the translocase domain. This gives these enzymes a high stability and high processivity on dsRNA. Our models predict that the Hel2i and RecA folds together confer similar stability on DNA for FANCM, but certain parts of Hel2i have further evolved to specifically engage the ATPase when bound to only branched DNA molecules.

By mutagenic screening across the Hel2i domain we identified K331, Y332, and H369 as key residues in defining both HJ binding and associated ATPase and branch migration activity. Importantly, subsequent AlphaFold 3 modelling of this region of FANCM engaged on a branched DNA substrate predicted that the same residues directly engage at the branchpoint. Thus, it is tempting to speculate that these residues act as pins, to pry open the DNA, in a mechanism similar to that used by helicases to open dsDNA. However, in our studies FANCM cannot unwind a static HJ, which aligns with previous studies which have universally failed to demonstrate helicase activity in FANCM (15,20,25,27). Instead, the reannealing of DNA adjacent to the breakpoint is necessary (a “mobile HJ”) for forward mobility of the enzyme, even though ATPase activity is spent upon binding to either static or mobile HJ. K331, Y332 and H369 are therefore critical couplers of junction binding and ATPase activity but not necessarily forward movement. This assures that branchpoint translocation only occurs on homologous sequences, and not across other structured DNA molecules that might form during replication or transcription.

Our analyses of the complexes formed between FANCM and HJ revealed that FANCM translocase domain can only bind HJ that already exists in the open planar conformation, a state that already has a low energetic barrier for branch migration. Other examples of branchpoint translocases that preferentially bind open conformation HJ include Rad54 (38) and *E*.*coli* RuvAB (39). Structural insights into both these proteins suggest they drive the conformational bias through dimeric interactions with the junction: binding to opposite arms and forcing a structural repositioning of the other two arms. However, in experiments using molar ratios of DNA and protein, we only observed single species in EMSA experiments when all junction is bound (Figure 6C), and we observe no binding when the HJ is in a stacked form. Therefore, unlike these other enzymes, FANCM is an opportunist. It binds only to four-way structures that are already in an open planar formation. One way that it might do this *in vivo* is by first binding to 3-way junctions, such as those found at stalled replication forks, and subsequently converting them into 4-way HJ that are exclusively planar. *In vitro* this has been observed as “replication fork reversal” by FANCM (17), and may be a way that restricts FANCM activity *in vivo* to certain substrates.

FANCM binding suppressed HJ cleavage by the GEN1 resolvase. This protection hints that FANCM’s translocase domain makes protective contacts with HJs, precluding GEN1 access and nuclease activity. Alternatively, maintaining the HJ in an open planar conformation could prevent GEN1 cleavage, as a previous study showed GEN1 requires the junction arms to be contorted at 90º out of plane for cleavage (40). Nuclease protection of replication forks and HJ is likely to be essential for maintenance of genome stability during branch migration of these substrates, but our results suggest FANCM could play a protective role on junctions even if it is not promoting their movement. In line with this, other studies suggest the nuclease XPF:ERCC1 could drive junction DNA cleavage in FANCM-deficient cells (9,20). XPF:ERCC1 more readily cleaves three-way replication/recombination intermediates accumulating in mitotic cells that lack FANCM and as such, four-way junction substrates for GEN1 may never accumulate. Further experiments should determine the roles of resolvases and other nucleases in cells in the absence of FANCM under various conditions, including in mitotic and meiotic settings.

ALT-positive cells represent a unique scenario. ALT is a cancer-specific process occurring independently of telomerase, and found in about 10% of cancers. ALT+ cancer cells have a specific dependence on FANCM. Because ALT relies primarily on break-induced replication for the copying of telomeric DNA during cell immortalisation, junction intermediates are prevalent due to elevated recombination, and require FANCM for their displacement (13). We found that the single-stranded DNA which accumulates at telomeres in FANCM-depleted ALT+ U2OS cells (12) cannot be rescued by the re-expression of FANCM-Hel2i mutants (Figure 7). This phenocopies the effects previously seen with ATPase dead FANCM or variants with defective Bloom syndrome complex binding (11,12). Our results add structure specific DNA binding to the list of required functions of FANCM in ALT suppression. Furthermore, consistent with FANCM limiting ALT by suppressing telomeric break-induced replication in G2, Hel2i mutants caused G2 cell cycle accumulation and loss of ALT cell viability. Identifying small molecules or inhibitors that block the Hel2i:DNA interaction could therefore have similar short-term anti-cancer properties as those proposed for FANCM-ATPase or FANCM-Bloom syndrome complex inhibitors.

FANCM deficiency is associated with cancer predisposition (7) and infertility disorders (41). Truncation mutations most strongly associate with these inherited phenotypes, but genome-wide association studies also link non-synonymous FANCM point mutations to breast cancer and reduced fertility (4). Many of the identified loss-of-function variants cluster in the N-terminal translocase domain and Hel2i region (42). Since single amino acid changes in the Hel2i domain led to complete loss of FANCM function in several assays, many naturally occurring mutants in this domain may also result in reduced or no FANCM activity. Structural models and assays developed here could stratify single amino acid variants in FANCM to improve prediction tools for these genetic disorders.

In summary, our results on the mechanism of structure specific binding by FANCM confirm both conserved and unique mechanisms of substrate engagement. While FANCM likely shares overall similarity with dsRNA binding enzymes, we identified unique residues in the Hel2i domain that determine its specificity for branched DNA substrates. The structure specific DNA binding properties of FANCM are likely to be further modified on particular DNA substrates by its associated proteins, including but likely not limited to MHF1 and MHF2(43), FAAP24(44), RMI1:RMI2 (45), hCLK2 (46) and PCNA(47). The overall stability and processivity of FANCM on branched DNA substrates have also made it an ideal landing pad for other DNA repair factors including the FA core complex and Bloom syndrome complex. Our data will have implications for understanding how and when these repair factors are activated for the overall maintenance of genome stability.

## Methods

### Protein models

To generate high-quality protein models for FANCM, a multi-step approach was employed. First, the FANCM sequence was submitted to Consurf(48), which provided a set of homologous sequences aligned with FANCM. This alignment was more robust than simply aligning FANCM with the HEF template, improving the quality of the subsequent homology model. The aligned sequence was then submitted to the SWISS-MODEL server for homology modelling. Among the numerous candidate templates (>200), the archaeal HEF protein had the most similar sequence to FANCM and was chosen as the primary template. To further enhance the sequence register, an alignment was generated using 80 vertebrate FANCM sequences along with HEF, and this improved alignment was then used to generate a homology model.

For modelling the isolated Hel2i domain of FANCM, the first 960 amino acids were submitted to the TrRosetta server(49), which generated a model using deep learning techniques. Additionally, the AlphaFold 2 model for FANCM was downloaded from the AlphaFold server(50). To generate AlphaFold 3 models of DNA-bound FANCM shown in Figure 5, the sequence of the translocase domain was uploaded, together with ATP and various structure-forming DNA sequences, to the AlphaFold server(51). The models are available in ModelArchive (modelarchive.org) with the accession codes ma-svk0u, ma-oj2j and ma-szcrz.

### Protein expression and purification

Full-length FANCM was purified as previously described(18). The genetic construct for expressing the N-terminally FLAG tagged FANCM translocase domain (residues 82-647), or mutant derivatives, was synthesised with a codon bias for expression in insect cells in a transposition-compatible vector pADC10(18). Plasmids were transformed into DH10^Multibac^ for integration into the MultiBac genome(52). Mutant forms were generated by *in vitro* mutagenesis, and expressed and purified using the same methods as wild-type protein. trFANCM and variants were expressed in Hi5 insect cells at 26ºC for 72 hours post infection in Sf900 Serum Free media. Cells were harvested by centrifugation at 1000 x g for 15 minutes (frozen and stored at –20ºC). All purification steps were conducted at 4 degrees C. Cells were lysed by sonication in buffer (50 mM HEPES pH 7.0, 300 mM NaCl, 5 mM DTT, 10% v/v glycerol, 1x mammalian protease inhibitor cocktail (APExBIO, K1007)). The lysate was clarified by centrifugation at 15000 x g for 30 minutes. Clarified lysate was applied to pre-equilibrated anti-FLAG M2 antibody resin (0.5-1 ml compact resin) and incubated with gentle rolling at 4 degrees for 1 hour. The resin was collected via centrifugation at 1000xg for 15 minutes at 4ºC, then washed 3x in the same manner with ∼30 ml purification buffer (50 mM HEPES pH 7.0, 300 mM NaCl, 5 mM DTT, 10% v/v glycerol). Resin was transferred to a gravity flow column, and an ATP wash (1 mM ATP, 2 mM MgCl^df^ in purification buffer) was performed. The ATP was washed away with 10x CV purification buffer, and protein was eluted with 4x CV of 0.2 mg/ml flag peptide in purification buffer.

GEN1 purification: pCDFDuet1-SUMO-GEN1_2-464_ was generated by gene synthesis, codon optimized for expression in E.coli. Plasmid was transformed into *E. coli* OverExpress C41(DE3) cells (Sigma). A single colony was selected and grown at 37ºC until the mid-log phase and induced overnight with 1 mM IPTG at 16ºC. Cells were harvested by centrifugation at 4000 rpm for 15 minutes at 4ºC, before the supernatant was discarded and the pellet was frozen at -80ºC. Pellets were resuspended in 30 ml GEN1 lysis buffer (1x PBS, 500 mM NaCl, 10% glycerol, 2 mM DTT, 1/100 PMSF, 1/100 bacterial protease inhibitor cocktail (Sigma-aldrich, ref: P8465-5ML), 10 mM imidazole) and lysed by sonication. The lysate was clarified by centrifugation at 15000 x g for 45 minutes at 4ºC. The clarified lysate was applied to pre-equilibrated nickel resin (1 ml compact resin) and incubated with gentle rolling at 4ºC for 1 hour. The resin was collected via centrifugation at 1000 x g for 15 minutes at 4ºC, then washed 3x in the same manner with 30 ml of purification buffer A (20 mM Tris pH 7.4, 500 mM NaCl, 10% glycerol, 2 mM DTT). An ATP wash with 2 mM ATP and 5 mM MgCl_2_ in purification buffer A was performed, before the resin was washed with 10x CV of purification buffer A. The resin was recovered via centrifugation at 1000 x g, for 15 minutes at 4ºC, then resuspended in 1 ml purification buffer A and transferred to a gravity flow column. Protein was eluted with 300 mM imidazole in purification buffer A. Eluted protein was dialysed into ULP1 buffer (25 mM Tris pH 7.4, 150 mM NaCl, 10% glycerol, 0.1% NP-40, 2 mM DTT) at 4ºC overnight, and the SUMO tag was cleaved using inhouse purified SUMO-Star protease. GEN1_2-464_ or the D30N nuclease dead protein was then dialysed into low salt buffer (20 mM MES pH 6.6, 75 mM NaCl, 10% glycerol, 2 mM DTT) overnight at 4ºC. Cation exchange chromatography was performed using a Cytiva Tricorn (5/50) column packed with Source 15S resin, with a linear gradient from 75 mM to 500 mM NaCl. Peak fractions were pooled, buffer exchanged into storage buffer (20 mM Tris pH 7.4, 100 mM NaCl, 0.1 mM EDTA, 2 mM TCEP, 5% glycerol), concentrated and flash frozen in liquid nitrogen, before the protein was stored at -80ºC.

MBP or MBP-FANCM-Hel2i (residues 293-447) was cloned by gene synthesis (Gene Universal) into pET16b and transformed into *E. coli* BL21(DE3) (Sigma). A single colony was selected and grown at 37ºC until the mid-log phase and induced overnight with 0.4 mM IPTG at 26ºC. Cells were harvested by centrifugation at 8000 xg for 15 minutes at 4ºC, before the supernatant was discarded and the pellet was frozen at -20ºC. Cells were lysed in buffer H (400 mM NaCl, 50 mM HEPES, 5 mM DTT, and 10% glycerol) and clarified by centrifugation at 16,100 x g for 1hr and incubated with ∼1 mL of pre-equilibrated compact amylose affinity resin beads (NEB). The resin was washed three times with buffer H. After the final wash, the resin was transferred to a gravity flow column and eluted in buffer H supplemented with 20 mM maltose. The most concentrated fractions were pooled (final concentration MBP 296.5 µM, MBP-FANCM-Hel2i 12μM) snap frozen in aliquots and stored at -80C.

### Generation of nucleic acid substrates

All nucleic acid substrates were generated by annealing synthesised ssDNA oligonucleotides (IDT). For static substrates used in EMSAs, all were generated as previously described(53). Asymmetric BXRH HJ structures were also generated by annealing, unlike previous protocols that relied on restriction digest of two arms (33,40). Asymmetric HJ structures were confirmed by EMSA in the presence and absence of Mg^2+^ as well as diagnostic restriction digest.

The migratable Holliday junction and replication forks were generated as previously described(20). Briefly, for 60 bp HJ, pairwise annealing of oligos IRDye700-X0m1 and X0m2, and Xm3 and Xm4, was performed. The annealed pairs were then incubated together for 1 hour at room temperature. For 50 bp HJ, pairwise annealing of oligos Xam1-3Dabcyl and Cy3-Xam2, and Xam3 and Xam4, was performed. For 60 bp migratable replication fork substrates, oligonucleotides IRDye700-X0M1 and X0M2.1/2, and FM and DS1, were pairwise annealed. The two pairs were then incubated together for 1 hour at room temperature. Glycerol (final concentration 5%) was added to the solution, before the annealed structures were purified as described for the 60 bp Holliday junction above.

### Enzyme-coupled ATPase assays

Reaction solutions containing 4 nM of FANCM translocase (WT, ΔHel2i, K331E, Y332A, H369E) or full-length FANCMc (WT), and 0, 2, 4, 8, 16, 32 and 64 nM of 30 bp non-migratable HJ were prepared in 1x ATPase buffer (20 mM Tris pH 8.0, 75 mM NaCl, 1 mM DTT) and 1x ATPase cocktail (0.2 mM NADH, 2 mM Phospho(enol)pyruvic acid tri(cyclo-hexylammonium) salt (sigma life science P7252-500MG, made up to 100 mM stock in milliQ water), 6 U/ml Pyruvate kinase (Roche, ref: 10128155001), 9 U/ml L-lactate dehydrogenase (Roche, ref: 10127230001). 1 mM of ATP and 2 mM of MgCl2 was added to the reaction solutions, before all solutions were transferred to a 384 well assay plate (Corning, black with clear flat bottom; ref: 3764). Reactions were incubated at 37ºC for 2 hours in a Perkin Elmer EnSpire Multimode Plate Reader. During this time, absorbance at 340 nm was monitored as a measure of NADH consumption over time. The reaction rate for the linear phase of each reaction was determined and a plot against [HJ] was made. Reactions were performed in triplicate and Michaelis-Menten enzyme kinetics analyses were performed using GraphPad Prism 10.

### Electromobility shift assays (EMSA)

For the translocase variants: 0, 0.25, 0.5, 1, 2, 4, 8 and 16 nM of FANCM translocase (WT, Hel2i, K331E, Y332A, H369E) were incubated with 0.25 nM of IRDye700 labelled 30 bp non-migratable Holliday junction or 60bp dsDNA in EMSA buffer (20 mM Tris pH 8.0, 300 mM NaCl, 10% glycerol, 0.005% NP-40) for 20 minutes at RT. For full-length FANCM: 0, 1, 2, 4, 8, 16, 32 and 64 nM of FANCM (WT, ΔHel2i) were incubated with 1 nM of IRDye700 labelled 30 bp non-migratable Holliday junction in EMSA buffer for 20 minutes at RT. Glycerol (additional 5%) was then added before reactions were resolved by electrophoresis through 6% PAGE gel in 1x TBE at a constant 70V for 1.5 hours. Results were imaged using the Li-Cor Odyssey CLx.

Asymmetric HJ EMSAs were conducted as above, with 2 nM asymmetric HJ and 2 nM translocase. Reactions run in the presence of Mg^2+^ had 0.5 mM MgCl_2_ in the binding reaction, native gel and running buffer. Reactions run in the absence of MgCl2 had 20 mM EDTA present in the binding reaction, native gel and running buffer.

For competition assays: 64 nM of FANCMc (WT, ΔHel2i) were incubated with 2 nM of non-migratable Holliday junction and 0, 2, 6, 18, 54, 162, and 486 nM unlabelled 60 bp dsDNA in EMSA buffer (20 mM Tris pH 8.0, 300 mM NaCl, 10% glycerol, 0.005% NP-40) for 20 minutes at room temperature to allow sufficient time for DNA binding. Glycerol (additional 5%) was then added to the reactions. Reactions were resolved by electrophoresis through 6% PAGE gel in 1x TBE at a constant 70V for 1.5 hours. Results were imaged using the Li-Cor Odyssey CLx.

10 nM of trFANCM was incubated with 2 nM of IRDye700 labelled 30 bp non-migratable Holliday junction and 0, 2, 6, 18, 54, 162, and 486 nM unlabelled 60 bp dsDNA in EMSA buffer (20 mM Tris pH 8.0, 300 mM NaCl, 10% glycerol, 0.005% NP-40) for 20 minutes at room temperature to allow sufficient time for DNA binding. Glycerol (additional 5%) was then added to the reactions. Reactions were resolved by electrophoresis through 6% PAGE gel in 1x TBE at a constant 70V for 1.5 hours. Results were imaged using the Li-Cor Odyssey CLx. All reactions were repeated at least 3 times, with similar results.

### Holliday junction branch migration assays

Full-length FANCM (0, 1, 2, 4, 8, 16, and 32 nM of WT, ΔHel2i) or trFANCM (0, 1, 4, and 16 nM of WT, ΔHel2i, RIG-I) were incubated with 20 nM of 50 bp migratable Holliday junction in reaction buffer (20 mM Tris pH 7.4, 75 mM NaCl, 5% v/v glycerol, 1 mM DTT, 0.1 mM EDTA, 0.005% NP-40, 1 mM MgCl_2_). 1 mM ATP was added to the reactions, which were then incubated at 37ºC for 10 minutes to initiate branch migration. The 10 μl reactions were deproteinized for 10 minutes at 37ºC with 3 mg/ml Proteinase K in branch migration loading dye (final concentrations 50 mM EDTA, 0.3% SDS, 10% glycerol, 0.17 mg/ml bromophenol blue). Reactions were resolved by electrophoresis through 6% polyacrylamide gel in 1x TBE at a constant 100V for 1 hour, then sybr gold post-stained for 20 minutes in 1xTBE. Results were imaged using the BioRad gel doc gel imager.

For the experiments with Hel2i mutants, reactions contained *16* nM of trFANCM (WT, ΔHel2i, RIG-I, D214A, L308Y, E309K, L321Y, R323D, K331E, Y332A, Q333A, I334Y, H369E, E372K) as above.

### Holliday junction and replication fork branch migration

Full-length FANCM vs FANCM translocase: 0, 1, 4, and 16 nM of FANCMc and trFANCM were incubated with 20 nM of IRDye700 labelled 60 bp replication fork, or 60 bp Holliday junction substrates, in reaction buffer (20 mM Tris pH 7.4, 75 mM NaCl, 5% v/v glycerol, 1 mM DTT, 0.1 mM EDTA, 0.005% NP-40, 1 mM MgCl_2_). 1 mM ATP was added to the reactions, which were then incubated at 37ºC for 10 minutes to initiate branch migration. The 10 μl reactions were deproteinized for 10 minutes at 37ºC with 3 mg/ml Proteinase K in branch migration loading dye (final concentrations 50 mM EDTA, 0.3% SDS, 10% glycerol, 0.17 mg/ml bromophenol blue). Reactions were resolved by electrophoresis through 6% polyacrylamide gel in 1x TBE at a constant 100V for 1 hour. Results were imaged using the Li-Cor Odyssey Clx.

### R-loop unwinding assays

To make plasmid-based R-loops 2µg of pUC19-Mu-switch-R-loop plasmid (Addgene: 134899) was incubated in a final reaction volume of 200µl containing 1x T7 polymerase reaction buffer (NEB), 25U T7 polymerase (NEB), 2.25mM each of CTP, GTP, ATP and 825nM ɑ-^32^P-UTP 3000 Ci/mmol (Perkin Elmer) for 1 hour at 37°C. The reaction was stopped by heat denaturation at 65°C for 20mins. 100µl of 1.05M NaCl, 30mM MgCl_2_ buffer was added to each reaction plus 2.5µg of Rnase A (EpiCentre) for 1hr at 37°C. R-loops where then purified as per (18), quantified using nanodrop and stored at 4°C.

R-loop branchpoint translocation reactions (10µl) contained 0.25nM of R-loop, 1mM ATP, 1, 3, 10 or 30nM FANCMc or FANCMcΔHel2i in R-loop buffer (6.6mM Tris pH7.5, 3% glycerol, 0.1mM EDTA, 1mM DTT, 0.5mM MgCl_2_) and incubated at 37°C for 10mins. Reactions were stopped by adding 2µl of stop buffer (10mg.ml^-1^ proteinase K (NEB), 1% SDS) and incubated for 15min at 37°C. 2µl of 50% glycerol was added to samples prior to loading onto 1% agarose TAE gels, run at 100V in TAE buffer (40mM Tris, 20mM acetic acid, 1mM EDTA) for 60 mins. Gels were then crushed between precut biodyene B membranes (Pall) for 1hour, exposed overnight to a GE phosphor-screen and imaged on a Typhoo scanner (GE Biosciences).

### GEN1 Holliday junction cleavage assays

IRDye700 labelled 60 bp non-migratable Holliday junction (10nM) was incubated with 0, 12.5, 25, 50, 100, 200, and 400 nM of trFANCM (WT, ΔHel2i, K331E, Y332A, H369E) in reaction buffer (20 mM Tris pH 7.4, 75 mM NaCl, 5% v/v glycerol, 1 mM DTT, 0.005% NP-40, 1 mM MgCl_2_) for 10 minutes at room temperature. 4 nM of GEN1_2-464_ was added to the reactions, which were then incubated at 37ºC for 5 minutes. The 10 μl reactions were deproteinized for 10 minutes at 37ºC with 3 mg/ml Proteinase K in branch migration loading dye (final concentrations 50 mM EDTA, 0.3% SDS, 10% glycerol, 0.17 mg/ml bromophenol blue). Reactions were resolved by electrophoresis through 6% polyacrylamide gel in 1x TBE at a constant 100V for 1 hour. Results were imaged using the Li-Cor Odyssey Clx.

### Differential scanning fluorimetry (DSF)

DSF was performed in a Roche LightCycler 480 II system. 10 µl reactions in LightCycler 480 Multiwell plates (white, 384 well) contained (20 mM Tris pH 7.4, 75 mM NaCl, 5% v/v glycerol, 1 mM DTT, 0.1 mM EDTA, 0.005% NP-40, 1 mM MgCl_2_), 0.1 µg trFANCM, 4x SYPRO orange, 2 µM bottom 1b oligonucleotide DNA (where indicated), and 1 mM ATP and 2 mM MgCl_2_ (where indicated). Plates were sealed using the manufacturer supplied sealing foils, then centrifuged at 1500 x g for 2 minutes. The instrument was programmed in the *Protein Melting* mode with a *SPYRO orange* detection format, following the manufacturer’s instructions. The instrument was warmed to 30ºC (4.8ºC/second), and reactions were held at 30ºC for 15 seconds, followed by an increase of 0.02ºC per second from 30ºC to 95ºC, before the instrument was cooled back to 30 ºC (2.5ºC/second) and reactions were held at 30ºC for 15 seconds. Plots for melting peaks were determined using the instruments *T*_*m*_ *calling* analysis mode. Both the raw data and melting peaks data was extracted in the text format. The raw data of arbitrary fluorescence against temperature (ºC) was analysed using the DSF world analysis software at https://gestwickilab.shinyapps.io/dsfworld/ (30) and T_m_s were determined. A plot of –(d/dT) Fluorescence against Temperature (ºC) was generated from the melting peaks data exported from the LightCycler 480 II.

### Circular dichroism

Samples were buffer exchanged into CD buffer (40 mM HEPES pH 8.0, 60 mM NaCl, 4 mM DTT, 10% glycerol) using desalting columns (company). CD spectra were measured at room temperature, using a 1 mm path length cuvette. Measurements were taken from 260-215 nm in a 0.1 nm step size in triplicate, using a Jasco spectropolarimeter. For an unstructured protein control, after measuring the native spectra, the wild-type FANCM translocase sample was heated to 80 degrees for 5 minutes before measuring CD again.

### Cell lines and culture

U2OS cells were obtained from CellBank Australia (STR profiled and mycoplasma free), and cultured in Dulbecco’s Modified Eagle’s Medium (DMEM, Gibco), supplemented with 10% fetal bovine serum (FBS, Sigma). Transfection mixes were diluted in Opti-MEM (Gibco).

### Plasmids and siRNA transfections

U2OS cells were reverse transfected with 25 nmol siRNA targeting the 3*′* UTR of FANCM (sequence: AAAGACCUCUCACAAUAUUtt from (25)) using lipofectamine RNAiMAX (ThermoFisher Scientitifc), according to manufacturer’s instructions. The next day, cells were transfected with 10 µg of empty vector, or wildtype or mutant FANCM constructs (pADC10-Flag-FANCM (18)), using FuGENE6 (Promega), according to manufacturer’s instructions. Cells were incubated for 2 days before being fixed for microscopy, or harvested for western blotting to check expression of FANCM. Clonogenic survival assays were performed as previously described (12).

### Microscopy

Cells grown on Alcian blue-treated coverslips (Sigma) were carefully washed in PBS before pre-extraction with KCM buffer for 10 mins (120 mM KCl, 20 mM NaCl, 10 mM Tris pH 7.5, 0.1% Triton X-100). Cells were washed in PBS, then fixed with 4% paraformaldehyde for 15 mins. Blocking was conducted for 1 hr in a solution containing 20 mM Tris–HCl pH 7.5, 2% (w/v) BSA, 0.2% (v/v) fish gelatin, 150 mM NaCl, 0.1% (v/v) Triton X-100, 0.1% (w/v) sodium azide and freshly added RNAse A at 100 µg/ml. After blocking, coverslips were stained with primary antibody against pRPA^ser33^ (Bethyl Laboratories, #A300-246A) at 1:500 dilution for 2 hrs at room temperature, and then with 1:500 dilution of anti-rabbit antibody conjugated to Alexa Fluor 568 for 1 hr (ThermoFisher Scientific, #A10042). Triplicates of PBST washes were performed after primary and secondary antibodies. After staining, another fixation step with 4% paraformaldehyde was conducted before coverslips were dehydrated using a graded ethanol series (75-100%). To detect telomeric DNA, coverslips were then mounted on glass slides with Tel C PNA FISH probes conjugated with Alexa Fluor 647 (HLB Panagene, #F2003), heated at 80oC for 3 min to induce denaturing, and hybridised overnight at 4oC. Cells were washed with PNA wash buffer A (70% formamide, 10 mM Tris pH 7.4) and then B (50 mM Tris pH 7.5, 150 mM NaCl and 0.08% Tween-20) for 15 mins each, before being rinsed in water. Coverslips were mounted in ProLongTM Gold antifade reagent containing DAPI (ThermoFisher Scientific) and imaged on the Zeiss Axio Imager microscope at 63x objective.

### Western blotting

Western blotting for FANCM expression was performed as previously described (54). Proteins were transferred onto PVDF membranes at 15 V overnight. Primary antibodies were mouse anti-FANCM clone CV5.1 (Novus Biologicals, #NBP2-50418, 1:1000 dilution) and anti-actin (Sigma, #a2066, 1:2000 dilution). Secondary antibodies were anti-mouse HRP (Agilent, #P0447, 1:10000 dilution) and anti-rabbit HRP (Agilent, cat #: P0448, 1:5000 dilution).

## Declaration of interest

AJD is a member of the scientific advisory board, and HAP a member of the board of directors of, Tessellate.bio. Conflicts are managed respectively by St Vincent’s Institute and Children’s Medical Research Institute. All other authors have no conflicts to declare.

## Acknowledgements

This research was performed on Wurundjeri and Gadigal land, funded by an Australian Government National Health and Medical Research Council grant (GNT1181110 to AJD and RBD) and a Medical Research Future Fund grant (MRFF2007488 to HAP). RBD was supported in part by a fellowship from the National Breast Cancer Foundation Australia. We acknowledge the support of the Victorian Government’s OIS Program.

