## Supplemental Figures 1-3 for "Mechanism of structure-specific DNA binding by the FANCM branchpoint translocase"

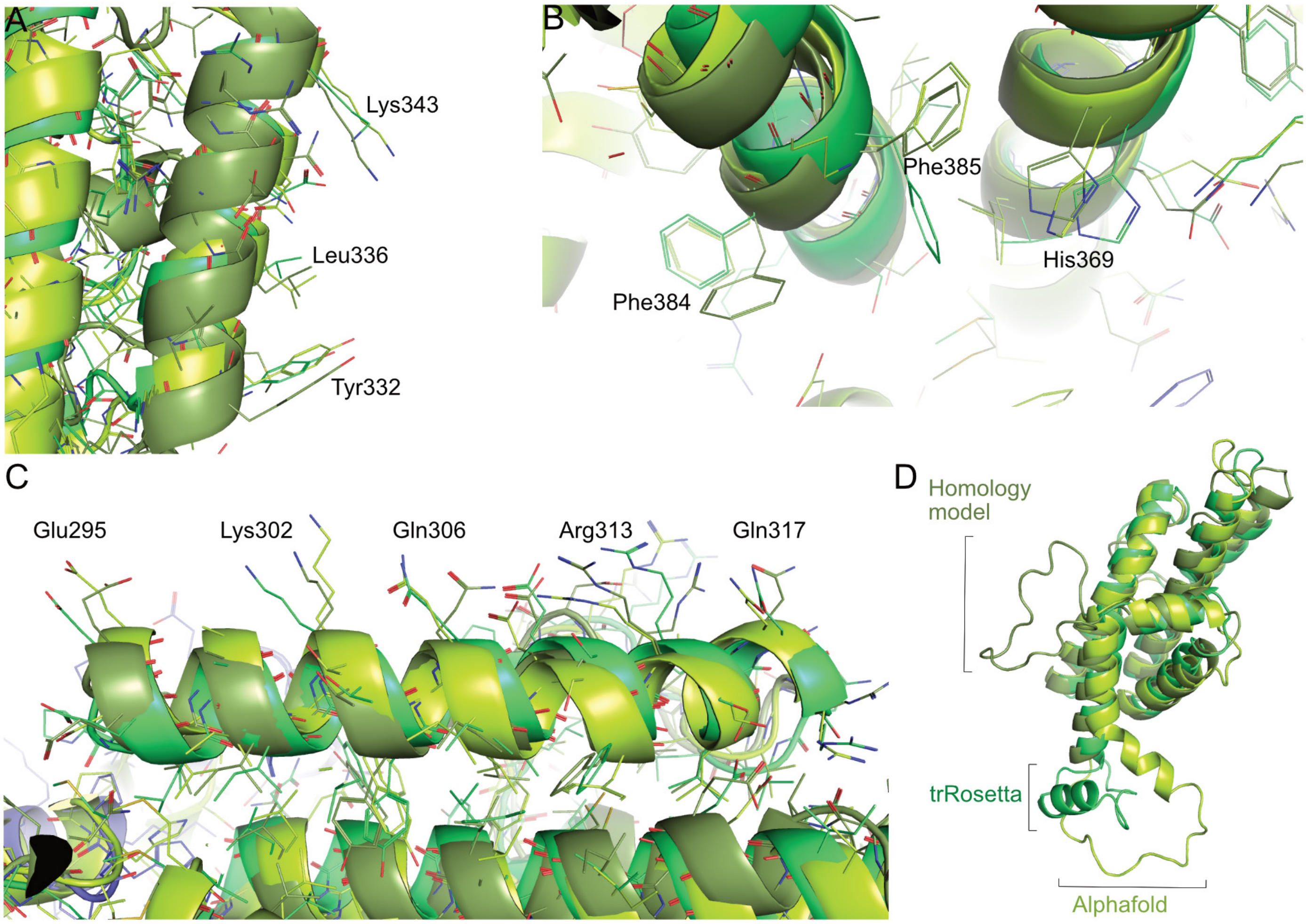


**Supplementary Figure 1: Comparison of the various FANCM Hel2i domain models.**
The TrRosetta, Alphafold and homology models were all superposed on their respective insert domain. A-C) Different regions of the insert domain highlighting that the three models share the same sequence register. D) The key region of divergence between the three models.

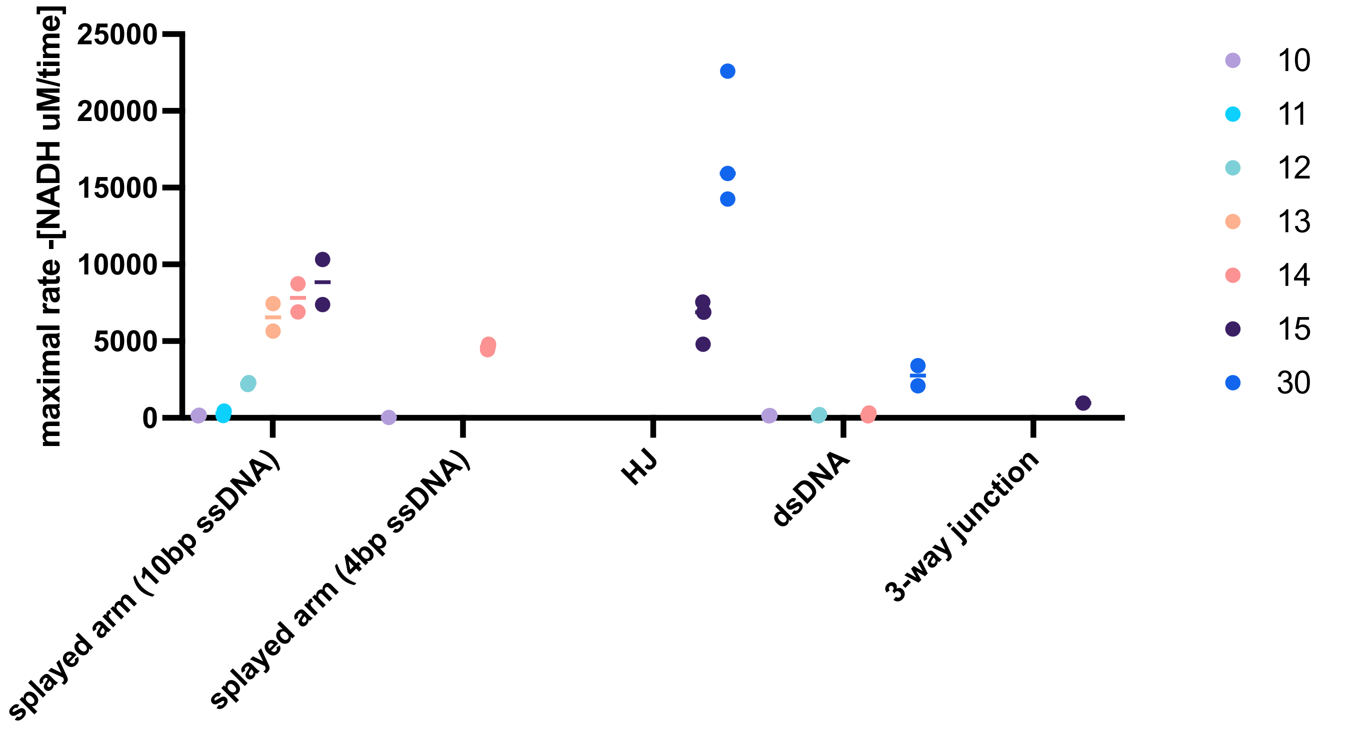


**Supplementary Figure 2. Stimulation of FANCM ATPase activity with different DNA structures**
ATPase assays were conducted as per materials and methods using 15nM FANCM and 30nM DNA substrate. Value represents the maximum rate during the linear phase of real-time ATPase assays, as measured by the depletion of NADH in the coupled reaction. Mean and individual values from n=3 replicates is shown. Different coloured labels represent different lengths of double-strand region. For splayed arm substrates, 10bp of 4bp ssDNA extensions were used.

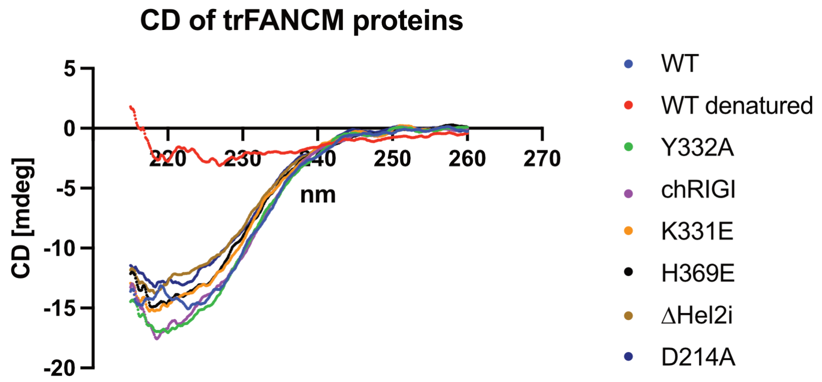


**Supplementary Figure 3: Circular dichroism of FANCM and various mutants used in the study**
CD was measured for all variants at 0.19 mg/mL. After the WT sample was measured, it was heated to 80°C for 5 mins and then re-measured as a denatured control.
